# Probing the fabric of the rare biosphere

**DOI:** 10.1101/020818

**Authors:** Bibiana G. Crespo, Philip J. Wallhead, Ramiro Logares, Carlos Pedrós-Alió

**Author notes:** Correspondence: Bibiana G. Crespo, Uni Research Environment, Center for Applied Biotechnology, Thormøhlens gate 49B, N-5006 Bergen, Norway. Present address: Uni Research Environment, Center for Applied Biotechnology, Thormøhlens gate 49B, N-5006 Bergen, Norway (Bibiana G. Crespo).

## Abstract

The relatively recent development of high-throughput sequencing (HTS) techniques has revealed a wealth of novel sequences found in very low abundance: the rare biosphere. Today, most studies of diversity of microorganisms are carried out almost exclusively with HTS techniques. However, culturing seems indispensable for diversity studies especially if the aim is exploring the rare biosphere. We have carried out a deep (1 million sequences per sample) pyrosequencing analysis of two marine bacterial samples and isolated a culture collection from one of them. We have shown that the collectors curves of the pyrosequencing data were close to reaching an asymptote. We have estimated the total richness of the samples and the sequencing effort necessary to obtain the total estimated richness experimentally. Comparing the pyrosequencing data and the isolates sequences we have found that isolation retrieves some of the rarest taxa and that the composition of rare taxa follows an annual succession. We have shown that increasing the number of tags sequenced would slowly uncover the isolated bacteria. However, even if the whole bacterial diversity was found by increasing the sampling depth, culturing would still be essential for the study of marine bacterial communities, especially if the target is the rare biosphere.

## Introduction

The question of how many species of living beings are there on Earth has intrigued ecologists and evolutionary scientists for decades (May, 1988; Erwin, 1991). One of the most recent estimates considered a number of 8.7 million species, but excluded bacteria and archaea from the estimate due to our ignorance of the microorganisms (Mora *et al.*, 2011). The International Census of Marine Microbes attempted to map the diversity of microbes in the oceans with novel high throughput sequencing techniques (Amaral-Zettler *et al*., 2010) but a global estimate was not attempted. Some estimates for marine bacterial species range from 10^4^ to 10^6^ based on different assumptions (Curtis *et al*., 2002; Hagström *et al*., 2002). Such a range of values, spanning several orders of magnitude, shows that we are far from a reasonable estimate.

Traditionally, bacteria were isolated in pure culture and then characterized biochemically and genetically until a new species could be formally described. It was realized that the bacteria able to grow in culture media were a small fraction of the bacterial cells that could be directly counted on a filter, a discrepancy named the “great plate count anomaly” (Staley & Konopka, 1985). Different studies estimated that only about 1% of the cells in natural waters could be cultivated (Pace, 1997; Eilers *et al*., 2000). Moreover, most of the cells in pure cultures were not the abundant ones in nature.

After the application of molecular cloning to natural systems (Giovannoni *et al*., 1990; Pace, 1997) a wealth of new taxa were found and, this time, they were the abundant ones in the oceans (DeLong, 1997; Pace, 1997). The drawback was that a sequence of the 16S rDNA did not provide much information about the physiology of the organism. Further, the realization that bacteria obtained in culture were mostly different from bacterial sequences obtained in clone libraries, produced what could be called the “great clone library anomaly”. Molecular methods could retrieve many sequences from the abundant organisms but missed the rare ones, and only occasionally a rare clone was found. Isolation, on the other hand, retrieved mostly rare bacteria and, occasionally an abundant one. This anomaly was a consequence of the fact that natural assemblages are formed a by a few taxa in very large concentrations and many taxa in very low concentrations. The problem can be easily visualized by looking at a rank-abundance curve (Pedrós-Alió, 2006). Primers for clone libraries will hybridize with the most abundant sequences over and over again before they find a rare target. Thus, only a fraction of the community will be available to cloning and sequencing. The relatively recent development of high-throughput sequencing (HTS) techniques and their application to natural microbial communities (Sogin *et al*., 2006) now provides an opportunity to solve the “great clone library anomaly”.

The study of microbial communities with such technologies has revealed a wealth of novel sequences found in very low abundance: the rare biosphere (Sogin *et al*., 2006). And different properties of the latter have been examined (Galand *et al*., 2009; Jones & Lennon, 2010; Pedrós-Alió, 2012; Caporaso *et al*., 2012; Lynch *et al*., 2012; Gibbons *et al*., 2013). Today, most studies of diversity of microorganisms are carried out almost exclusively with such HTS techniques. Yet, culturing seems indispensable for diversity studies (Donachie *et al*., 2007; Shade *et al*., 2012; Lekunberri *et al*., 2014), especially if the aim is exploring the rare biosphere. In this regard, Shade *et al*. (2012) compared the outputs of a shallow (~ 2 000 sequences per sample) pyrosequencing analysis of the bacteria collected from a soil sample and the isolates cultured from the same sample. They found that 61% of the cultured bacteria were not present in the pyrosequencing dataset, demonstrating that culturing provided a fruitful route to the rare biosphere that was complementary to sequencing. These authors postulated that they would have found that remaining cultured bacteria in the pyrosequencing dataset if hey had increased the sequencing depth.

But the question “can all the bacterial taxa isolated in culture from a sample be found by HTS?” has not been addressed in a direct and sufficiently exhaustive way. This question has several implications. First, an in depth comparison of both cultures and sequences should eventually solve the “great clone library anomaly”. Second, it should provide an estimate of the total number of bacterial taxa in a sample. And third, it would allow a calculation of the sequencing effort necessary to retrieve all the diversity in a sample. We have addressed this question by carrying out a deep (1 million sequences per sample) pyrosequencing analysis of two marine bacterial samples and isolating a culture collection from one of them.

Comparison of both data sets allowed us to bracket the dimensions of the rare biosphere, at least in these two marine samples.

## Material and methods

### 1. Study area and sampling

Samples were taken in the NW Mediterranean Sea during cruise SUMMER between 13^th^ and 22^nd^ of September 2011, on board the RV “García del Cid”. Specifically, the samples for this study were collected at Station D, an open sea station at 40°52’N and 02°47’E (Table 1, and Pedrós-Alió *et al*., 1999). The surface sample was taken at 5 m on 15^th^ September and the bottom sample was collected at 2 000 m depth on 17^th^ September.

**Table 1.** Summary of location and depth (m) of samples, total sequences before (Raw Tags) and after (Final Tags) cleaning, richness (S) computed as total Operational Taxonomic Units (OTUs) clustered at 97% identity, percentage of singletons. Diversity estimates according to the Shannon diversity index (H’), Simpson diversity (D) and Pielou’s evenness (J). Richness (S) estimates according the non-parametric methods Chao Index (SChao) and Abundancebase Coverage Estimator (ACE) (S_ACE_) and the fitting of the accumulation curve to a Weibull Cumulative three parameter function (Sw_eibullFit_) which model efficiency was 99% for the surface and the bottom samples fitting (see Fig. 1). Estimates of the sampling effort (number of Tags) that would be needed to obtained the 99% and 99.9% of the total richness estimated by the Chao Index (S_Chao_) and the Weibull cumulative function fitting (S_WeibullFit_).

|  | Surface | Bottom |
| --- | --- | --- |
| Lat, Long | 40°52'N, 02°47'E | 40°52'N, 02°47'E |
| Depth (m) | 5 | 2 000 |
| Raw Tags | 713 076 | 970 346 |
| Final Tags | 500 262 | 574 960 |
| OTUs 97 % identity (S) | 1 400 | 4 460 |
| Singletons (% OTUs) | 17.86 | 17.20 |
| Diversity estimates: |  |  |
| $H'$ (Shannon diversity index) | 3.26 | 4.75 |
| D (Simpson diversity) | 0.45 | 0.66 |
| $J'$ (Pielou's evenness) | 0.45 | 0.57 |
| Richness estimates (estimated S): |  |  |
| Chao Index ($S_{Chao}$) | 1 643 | 5 029 |
| ACE ($S_{ACE}$) | 1 556 | 4 924 |
| Weibull Cumulative Fitting ($S_{WeibullFit}$) | 1 519 | 4 946 |
| Sampling effort estimates (Number of Tags) for: |  |  |
| 99 % of $S_{Chao}$ | $1.33 \times 10^6$ | $1.04 \times 10^6$ |
| 99.9 % of $S_{Chao}$ | $2.47 \times 10^6$ | $2.03 \times 10^6$ |
| 99 % of $S_{WeibullFit}$ | $1.92 \times 10^6$ | $1.96 \times 10^6$ |
| 99.9 % of $S_{WeibullFit}$ | $4.02 \times 10^6$ | $3.91 \times 10^6$ |

Sampling was done with Niskin bottles mounted on a rosette with a conductivity-temperature-depth (CTD) profiler. Water was prefiltered through a 200 μm mesh and immediately processed on board. To collect microbial biomass, between 5 and 15 L of sea-water were prefiltered through a 3 μm pore size Durapore filter (Millipore, Cork, Ireland) and free-living bacterial biomass was collected on a 0.22 μm pore size Sterivex filter (Durapore, Millipore). The filtration was done in succession using a peristaltic pump. The 0.22 μm pore size Sterivex unit was filled with 1.8 ml of lysis buffer (40 mM EDTA, 50 mM Tris-HCl, 0.75 M sucrose) and stored at −80°C. DNA was extracted by a standard protocol using phenol/chloroform (details in Schauer *et al*., 2003).

### 2. 454-pyrosequencing and noise removal

Purified DNA samples were submitted to the Research and Testing Laboratory (Lubbock, Texas, USA) for amplification of the 16S rRNA gene. Tag-pyrosequencing was done with Roche 454 Titanium platform following manufacturer protocols (454 Life Science). Primers 28F (5’ -GAGTTTGATCNTGGCTCAG) and 519R (5’ -GTNTTACNGCGGCKGCTG) were used for amplification of the hypervariable regions V1-V3; approximately 400 bp long tags were obtained. PCR and subsequent sequencing are described in Dowd *et al*. (2008). 713 076 and 970 346 tags were retrieved from the surface and the bottom samples, respectively (Table 1). These data has been deposited in EMBL with accession number PRJEB9061.

The raw tag-sequences were processed using QIIME (Caporaso *et al*., 2010). Briefly, to reduce sequencing errors and their effects, the multiplexed reads were first trimmed, quality-filtered and assigned to the samples, surface or bottom. The filtering criteria included a perfect match to the sequence barcode and primer, at least 400 bp in length, a quality score window of 50 bp and a minimum average quality score of 28. Additionally, denoising was used to reduce the amount of erroneous sequences (Quince *et al*., 2011). The final number of tags was reduced after this processing to 500 262 for the surface sample and to 574 960 for the bottom sample (Table 1). The sequences were then clustered into Operational Taxonomic Units (OTUs) based on the relatedness of the sequences (97% similarity) with UCLUST. Afterwards, a representative sequence from each OTU was selected. To identify potential chimera sequences, the dataset was subjected to the ChimeraSlayer implemented in Mothur (Schloss *et al*., 2011). Then, taxonomy assignment was made with QIIME by searching the representative sequences of each OTU against the SILVA 16S/18S rDNA non-redundant reference dataset (SSU Ref 108 NR) (Quast *et al*., 2013) using the Basic Local Alignment Search Tool (BLAST) and an e-value of 0.03. Chimera, chloroplast, eukarya and archaea sequences were removed from the output fasta file that was used for building a table with the OTU abundance of each sample and the taxonomic assignments for each OTU.

In addition to the sequences obtained in the present study, we used a 454 pyrosequencing dataset from cruise MODIVUS, for comparative purposes. Cruise MODIVUS took place in September 2007 and sampled a transect between the Blanes Bay Microbial Observatory (BBMO, http://www.icm.csic.es/bio/projects/icmicrobis/bbmo/), a coastal sampling site north of Barcelona (41°40’N, 02°48’E) and Station D (40°52’N, 02°47’E), the open sea station sampled here. The samples used for pyrosequencing included surface samples along the transect and a vertical profile down to 2000 m at Station D. The MODIVUS data have been published in Pommier *et al*. (2010) and Crespo *et al*. (2013). Due to the limitations of the technique at the time, these sequences only included de V3 region of the 16S rDNA (~68 bp).

### 3. Isolation of bacterial cultures

Isolates were obtained on board by plating 100 μl of undiluted and 10x diluted sea-water from the surface sample, in triplicates, onto modified Zobell agar plates (i.e. 5 g peptone, 1 g yeast extract and 15 g agar in 1 l of 0.2 μm filtered 75% sea water). Agar plates were incubated at *in situ* temperature (~20 °C), in the dark, for 14 days. 326 bacterial colonies were selected and the cultures were subsequently purified by re-isolation three times in a month. Next, isolates were grown at 20 °C on the same liquid medium and stored at −80 °C with 25% (v/v) glycerol. 200 μl of these cultures were place in 96 well plates, diluted 1:4 and heated (95°C, 10 min) to cause cell lysis, so available DNA could be used as a template in Polimerase Chain Reactions (PCR). PCR, using Taq polymerase (Boehringer-Mannheim), of the Internal Transcribed Spacer (ITS) were done to select as many different species as possible from the 326 isolates. ITS length is species specific and therefore allows to differentiate the isolates (Fisher & Triplett 1999; Scheinert *et al*. 1996). ITS amplification was done using primers ITS-F (5’-GTCGTAACAAGGTAGCCGTA) and ITS-R (5’-GCCAAGGCATCCACC) and the following thermal conditions: 94°C for 2 min, then 32 cycles of 94°C for 15 sec, 55°C for 30 sec, 72°C for 3 min, followed by one cycle of 72°C for 4 min and 4°C on hold. According to their different ITS patterns, 148 isolates were chosen out of 326, including some replicates, and their 16S rRNA gene were then amplified using bacterial 16S rRNA gene primers 27F (5’-AGAGTTTGATCMTGGCTCAG) and 1492R (5’-GTTTACCTTGTTACGACTT). The thermal conditions were as follows: 94°C for 5 min, then 30 cycles of 94°C for 1 min, 55°C for 1 min, 72°C for 2 min, followed by one cycle of 72°C for 10 min and 4°C on hold. Nearly the full-length 16S rRNA gene (aprox. 1 300 bp) was sequenced in GENOSCREEN (Lille Cedex, France). Taxonomical assignment was done by BLAST searches in the National Center for Biotechnology Information (NCBI) website. The 16S rRNA sequences have been deposited in EMBL with accession numbers LN845965 to LN846112.

In addition to the SUMMER culture collection obtained for the present study, we also used three previously isolated collections for comparison. The isolation procedures were the same as above, but the samples were collected at the Blanes Bay Microbial Observatory (BBMO). Station D sampled in the present work is about 100 km offshore from the BBMO. Culture collection BBMO-1 consisted of over 300 isolates collected between 2001 and 2004 at different times of the year. Collections BBMO-2 and BBMO-3 were smaller collections isolated in February and September 2007 respectively. Since these cultures were isolated and used for different purposes, many of them had only partial sequences of the 16S rDNA. A description of these collections was published in Lekunberri *et al*. (2014).

### 4. Richness, sampling effort estimates, and diversity of 454 pyrosequecing data

Richness (S) was computed as the total number of OTUs (97% similarity) in each sample. Estimates of total richness were calculated using two packages of the free software R (R Core Team, 2013). The “vegan” package (Oksanen *et al*., 2013) was used for non-parametric (S_Chao_, Abundance-based Coverage Estimator [ACE]) estimations. The “drc” package (Ritz & Streibig, 2005) was used to estimate richness by fitting the species accumulation curves to a mathematical function. We fitted several mathematical functions to our data (Flather, 1996). The best fits were given by Michaelis-Menten, Rational, and Weibull Cumulative functions. The three parameter Weibull Cumulative function was the best of all according to the coefficient of determination (R^2^), the residual sum of squares (RSS), and Akaike’s information criterion (AIC) (Supplementary Table 1). The Weibull cumulative function takes the form: 

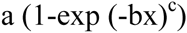
where a, b and c are fitted coefficients, and a is the maximal number of species predicted by the model, i.e. the asymptote.

The sampling effort necessary to obtained 99% and 99.9% of total estimated richness was calculated from the extrapolation of the species accumulation curves by fitting the Weibull Cumulative function and from the non-parametric estimation of Chao 1 using the method described in Chao *et al*. (2009).

Shannon (H’) and Simpson (D) diversity indexes were used to calculate diversity: 

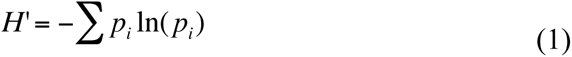

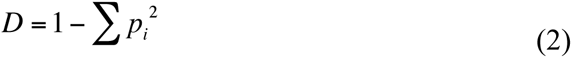
where *p*_i=_ N_i_/N, the number of individuals of species *i* divided by the total number of individuals in the sample (N). Finally, evenness was computed with the Pielou index: 

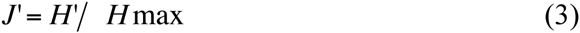
where H’ is the Shannon diversity index and H_max_ is the maximal possible Shannon diversity index if all the species were equally abundant: 

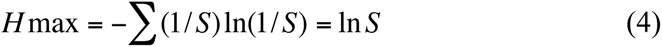
where S is the total number of OTUs (richness). Diversity and evenness of each sample were calculated using the “vegan” package (Oksanen *et al*., 2013).

Rank-abundance plots of the isolated cultures and the 454 pyrosequencing data were done using the “BiodiversityR” package (Kindt & Coe, 2005) and the accumulation curves using the “vegan” package (Oksanen *et al*., 2013) of R (R Core Team, 2013).

### 5. Comparison of 454-pyrosequencing tags and isolates

Comparison between isolates and 454 tag-sequences was done running BLASTn locally. Thus, the isolate sequences were searched for in the 454 tag-sequence datasets and vice versa, and only the reciprocal matches between these two searches were considered. The output was filtered using R (R Core Team, 2013) according to a 99% of identical nucleotide matches, 75%-100% of coverage of the isolate sequence and a bit-score higher than 100. In all the cases the e-value was lower than 0.0001.

Since the primers used for Sanger sequencing of the isolates and those used for the pyrosequencing of the environmental DNA were different, the possibility existed of different biases that could prevent detection of the cultures in the 454 dataset. Multiple alignments of the sequences of the isolates and the sequences of the primers used in the pyrosequencing analysis were done using the software Genious. The multiple alignments were used to check that the 454 primers hybridized with the sequences of all the isolates.

In order to estimate the sequencing effort necessary to find all the isolates in the sequence dataset, we did a kind of “rarefaction” analysis with detected isolates in the Y-axis and number of 454 sequences or number of OTUs in the X-axis. The 454 tag-sequences were subsampled in 5% intervals 1000 times and the percentage of coincidence with the isolate sequences was calculated for each new 454 tag-sequence dataset and plotted against the corresponding number of sequence tags analyzed. The same was done with the number of OTUs. Calculations were done using R (R Core Team, 2013).

## Results

### 1. Pyrosequencing dataset

Richness (S), computed as the total number of OTUs, was higher in the bottom (4 460) than in the surface (1 400) sample (Table 1). In both samples only ~17% of the OTUs were singletons (an OTU represented by a single sequence) (Table 1). Evenness (J’) and diversity (H’ and D) were also higher in the bottom than in the surface sample (Table 1). Richness estimations calculated with non-parametric methods, Chao Index and ACE, and with parametric methods (Figure 1) were slightly higher than the actual number of OTUs (S) observed as expected (Table 1). None of the estimates were significantly different from each other (z-tests on the differences, all P > 0.05). The mathematical function that best fitted the species accumulation curves was the Weibull Cumulative function (99% model efficiency or R^2^ =0.99) (Figure 1). Fitting the species accumulation curve allowed both calculation of the total richness of the samples by extrapolation (as explained above), and estimation of the sequencing effort necessary to obtain the total estimated richness experimentally (Table 1). Our estimates indicated that the sequencing effort necessary to retrieve 99% and 99.9% of the total estimated richness should be 4 to 8 times higher that the one applied in this study (Table 1). These numbers are approximately twice higher that the estimations obtained from the Chao index (Chao *et al*., 2009) (Table 1).

**Figure 1.**
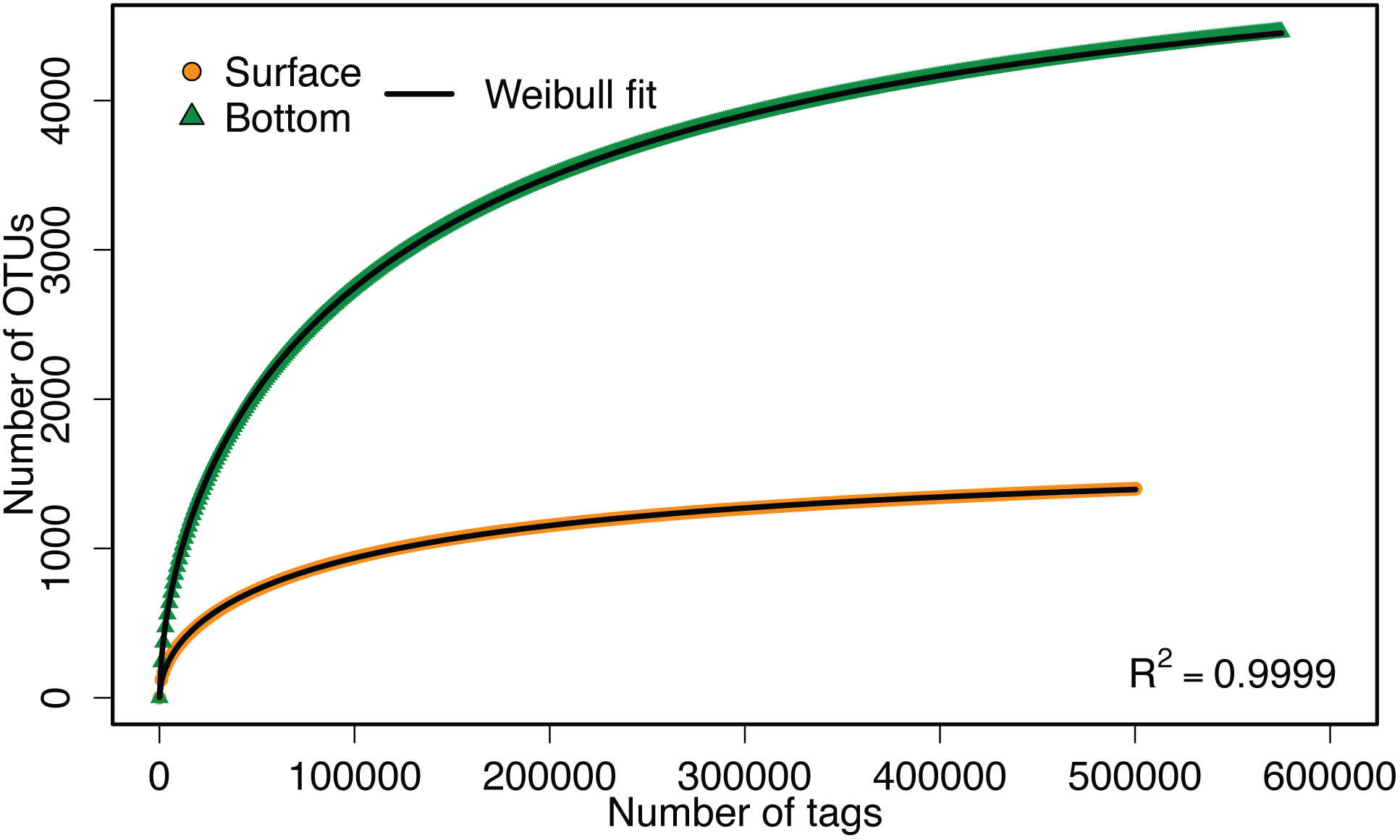
OTU accumulation curves of the surface (orange circles) and bottom (green triangles) samples. The black line is the Weibull cumulative function fit with model efficiency of 99% (R^2^=0.99).

Rank-abundance curves (Figure 2) showed that the bacterial assemblages from both samples were characterized by few abundant and many rare OTUs. The most abundant OTU was more abundant in the surface than in the bottom sample, in agreement with the lower evenness found for the surface sample (Table 1). The abundance of the most abundant OTU in the bottom sample was close to the abundance of the second most abundant OTU in the surface sample.

**Figure 2.**
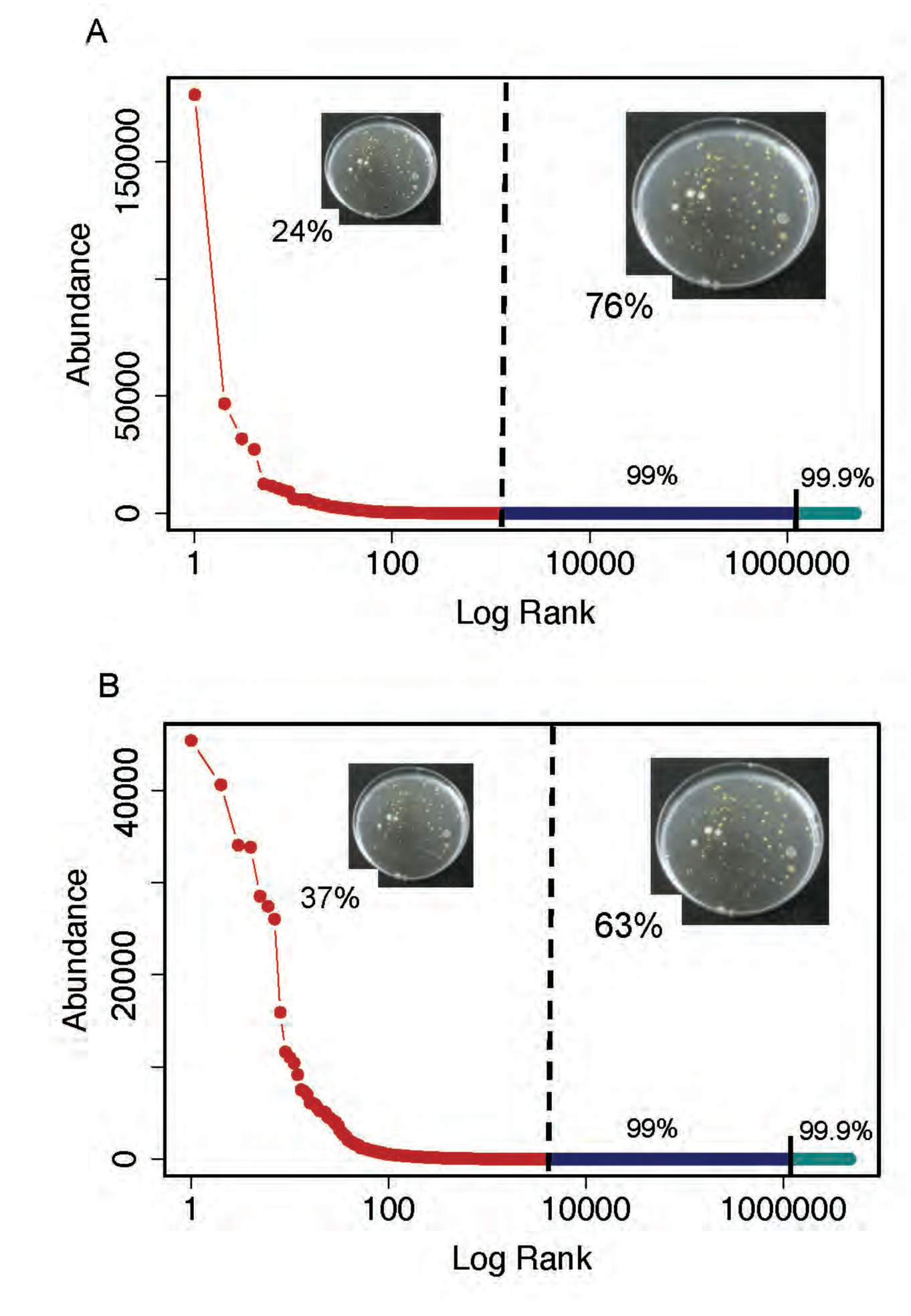
Rank-abundance plots of surface (A) and bottom (B) samples. The red line is the rank-abundance plot calculated with the actual data. The blue lines show the estimates of the sampling effort necessary to retrieve 99% (dark blue) or 99.9% (light blue) of the total estimated richness calculated by extrapolation of the species accumulation curve using the Weibull cumulative function. The vertical black line separates the real data (left) from the estimates (right). The percentage of cultured isolates found in the 454-pyrosequencing datasets is indicated at the left side of the black vertical line. The percentage of cultured isolates not found in the 454-pyrosequencing datasets, and that would presumably be found by increasing the sampling effort, is indicated at the right of the black vertical line. Insert pictures show some of the bacterial cultures grown from the surface sample. Font size and pictures are scaled according to the percentage of cultured isolates found or not found in the 454-pyrosequencing datasets.

### 2. Culture collection

Bacterial isolation from the sample collected at the surface retrieved 148 cultures belonging to 38 different species. The most frequent bacterium in the collection was *Erythrobacter citreus*, isolated 37 times, while 17 species were isolated only once. A rank abundance plot of the 38 species can be seen in Fig. 3. The isolates belonged to the phyla *Actinobacteria* (4 isolates), *Bacteroidetes* (4 isolates) and *Firmicutes* (2 isolates) and to the *Proteobacteria* classes *Alpha-proteobacteria* (18 isolates) and *Gamma-proteobacteria* (10 isolates). The names of all the isolates are shown in Table 2 and supplementary Table 2.

**Figure 3.**
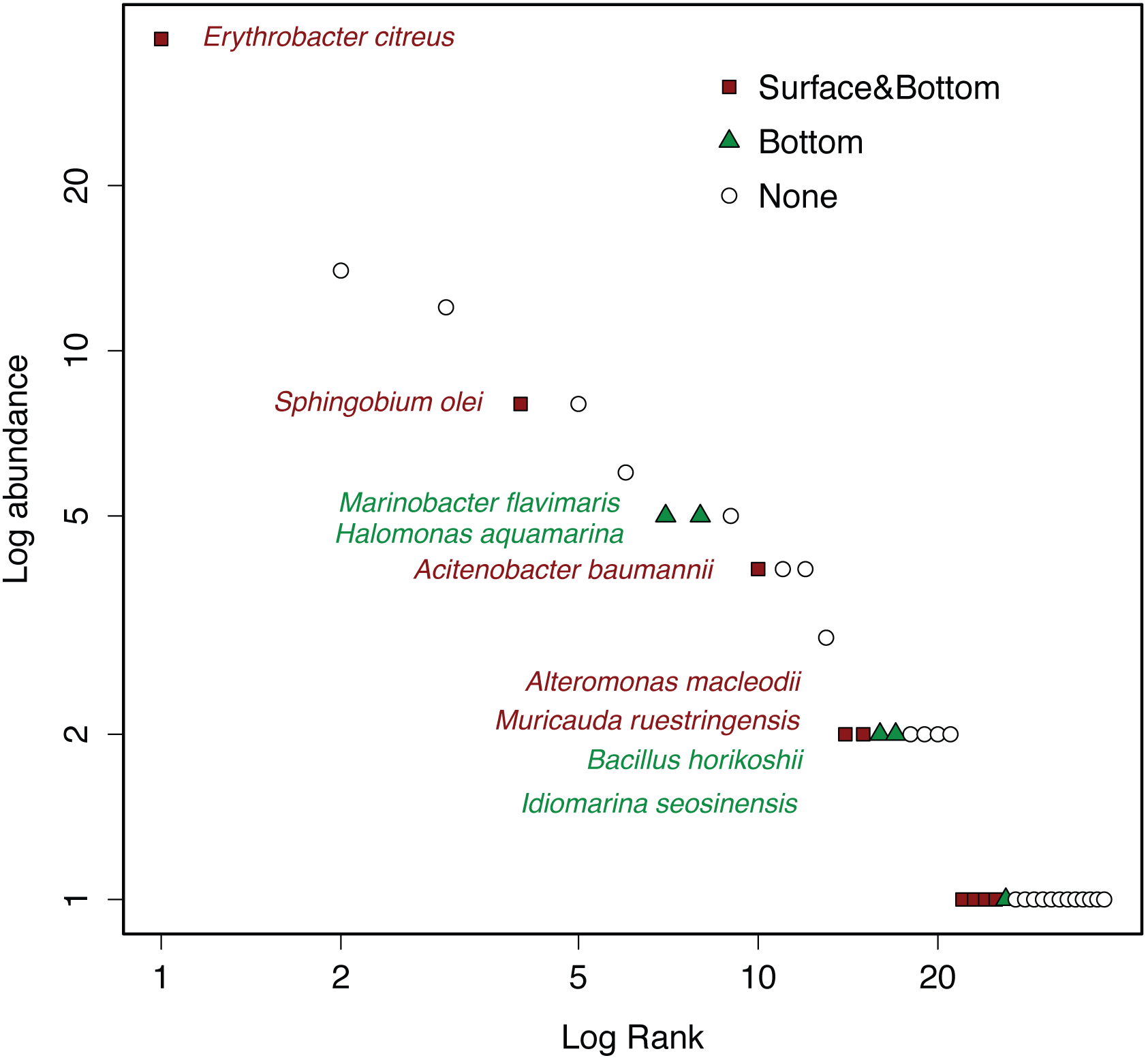
Rank-abundance plot of the 38 isolated bacterial species. The maroon squares indicate the cultured isolates found in both the surface and bottom 454-pyrosequencing datasets, the green triangles indicate the cultures isolated found only in the bottom 454-pyrosequencing dataset, and the white circles indicate the cultures that were not found in any of the 454-pyrosequencing datasets. A list of the isolated bacterial species can be found in Table 2 and supplementary Table 2.

**Table 2.** Isolates’ closest relative according to BLAST results, *%* of identity with the BLAST reference strain (identity BLAST), GenBank accession number of the BLAST reference strain, number of tags matching the isolates sequences in the surface and bottom samples (Tags in Surface, Tags in Bottom), percentage of the tags in the surface and bottom samples (% Surface, % Bottom) and number of isolates of each taxa sequenced. *Actino* (*Actinobacteria*)*, Bact* (*Bacteroidetes*)*, Firm* (*Firmicutes*), Alpha-*P* (Alpha-*Proteobacetria*) and Gamma-*P* (Gamma-*Proteobacteria*).

| Isolates' closest relative | Identity BLAST | GenBank accession number | Tags in Surface | % Surface | Tags in Bottom | % Bottom | Isolates number |
| --- | --- | --- | --- | --- | --- | --- | --- |
| *Uncultured Brevundimonas* sp. (Alpha-P) | 99.9 % | JX047099 | 76 | 1.52x10<sup>-2</sup> | 172 | 2.99x10<sup>-2</sup> | 1 |
| *Alteromonas macleodii* str. 'Balearic Sea AD45' (Gamma-P) | 100 % | CP003873 | 40 | 8.00x10<sup>-3</sup> | 7526 | 1.31 | 2 |
| *Sphingobium olei* (Alpha-P) | 100 % | HQ398416 | 34 | 6.80x10<sup>-3</sup> | 232 | 4.04x10<sup>-2</sup> | 8 |
| *Erythrobacter citreus* (Alpha-P) | 100 % | EU440970 | 31 | 6.20x10<sup>-3</sup> | 861 | 1.50x10<sup>-1</sup> | 37 |
| *Citromicrobium* sp.(Alpha-P) | 100 % | HQ871848 | 22 | 4.40x10<sup>-3</sup> | 39 | 6.78x10<sup>-3</sup> | 1 |
| *Acinetobacter baumannii* (Gamma-P) | 100 % | JX966437 | 16 | 3.20x10<sup>-3</sup> | 128 | 2.23x10<sup>-2</sup> | 4 |
| *Bizionia* sp. (*Bact*) | 100 % | EU143366 | 13 | 2.60x10<sup>-3</sup> | 66 | 1.15x10<sup>-2</sup> | 1 |
| *Muricauda ruestringensis* (*Bact*) | 99 % | JN791391 | 4 | 8.00x10<sup>-4</sup> | 92 | 1.60x10<sup>-2</sup> | 2 |
| *Microbacterium jejuense* (*Actino*) | 100 % | AM778450 | 1 | 2.00x10<sup>-4</sup> | 15 | 2.61x10<sup>-3</sup> | 1 |
| *Marinobacter flavimaris* (Gamma-P) | 100 % | AB617558 | 0 | 0 | 174 | 3.03x10<sup>-2</sup> | 5 |
| *Bacillus* sp. (*Firm*) | 100 % | AM950311 | 0 | 0 | 17 | 2.96x10<sup>-3</sup> | 1 |
| *Bacillus horikoshii* (*Firm*) | 100 % | JQ904719 | 0 | 0 | 8 | 1.39x10<sup>-3</sup> | 2 |
| *Halomonas aquamarina* (Gamma-P) | 100 % | AB681582 | 0 | 0 | 1 | 1.74x10<sup>-4</sup> | 5 |
| *Idiomarina seosinensis* (Gamma-P) | 99.9 % | EU440964 | 0 | 0 | 1 | 1.74x10<sup>-4</sup> | 2 |

### 3. Comparison of isolates and sequences

Only 14 (37%) of the 38 different species were found in the 454 tag-sequence datasets: one *Actinobacteria*, two *Bacteroidetes*, two *Firmicutes*, four Alpha-*proteobacteria* and five *Gamma-proteobacteria* isolates (Figure 3, Table 2). Surprisingly, the number of cultures found in the 454 tag-sequence dataset was higher in the sample collected at 2 000 m (37%) than in the surface sample (24%), even though the latter was the sample used for isolation of the bacterial cultures (Figure 3, Table 2). Nine species were found in the sequences from both samples (maroon in Figure 3), 5 were found in the bottom sample only (green in Figure 3) and 24 were not found in either sample (empty symbols in Figure 3). Practically all the 454 tag-sequences that matched the sequences from the isolates belonged to rare OTUs (<1% of the total tags). Only the OTU matching the isolate *Alteromonas macleodii* str. ‘Balearic Sea AD45’ (*Gamma-Proteobacteria*) was somewhat abundant (1.3%) in the bottom sample (Table 2). Further, all the matching sequences made a larger percentage of the assemblage at the bottom sample than at the surface sample.

In order to examine the rate of appearance of cultured species in the sequence database, we calculated a kind of “rarefaction curves”, by randomly resampling either the OTUs or the 454-pyrosequecing tags in 5% increases, and checking how many cultured species had been found in each case (Figure 4). These curves showed that the rate of appearance of the isolates in the 454 tags datasets as the sequencing depth increased was slower for tags than for OTUs but in neither case was an asymptote reached (Figure 4). The rate of increase of retrieved isolates versus OTUs was constant, but the rates were different at the two depths: one new isolate was retrieved every 50 OTUs in the surface sample and every 117 OTUs in the bottom sample (Figures 4A-B). The increase in retrieved isolates with respect to the number of tags showed a nonlinear response, implying that the sampling effort had to increase as the number of isolates increased (Figures 4C-D).

**Figure 4.**
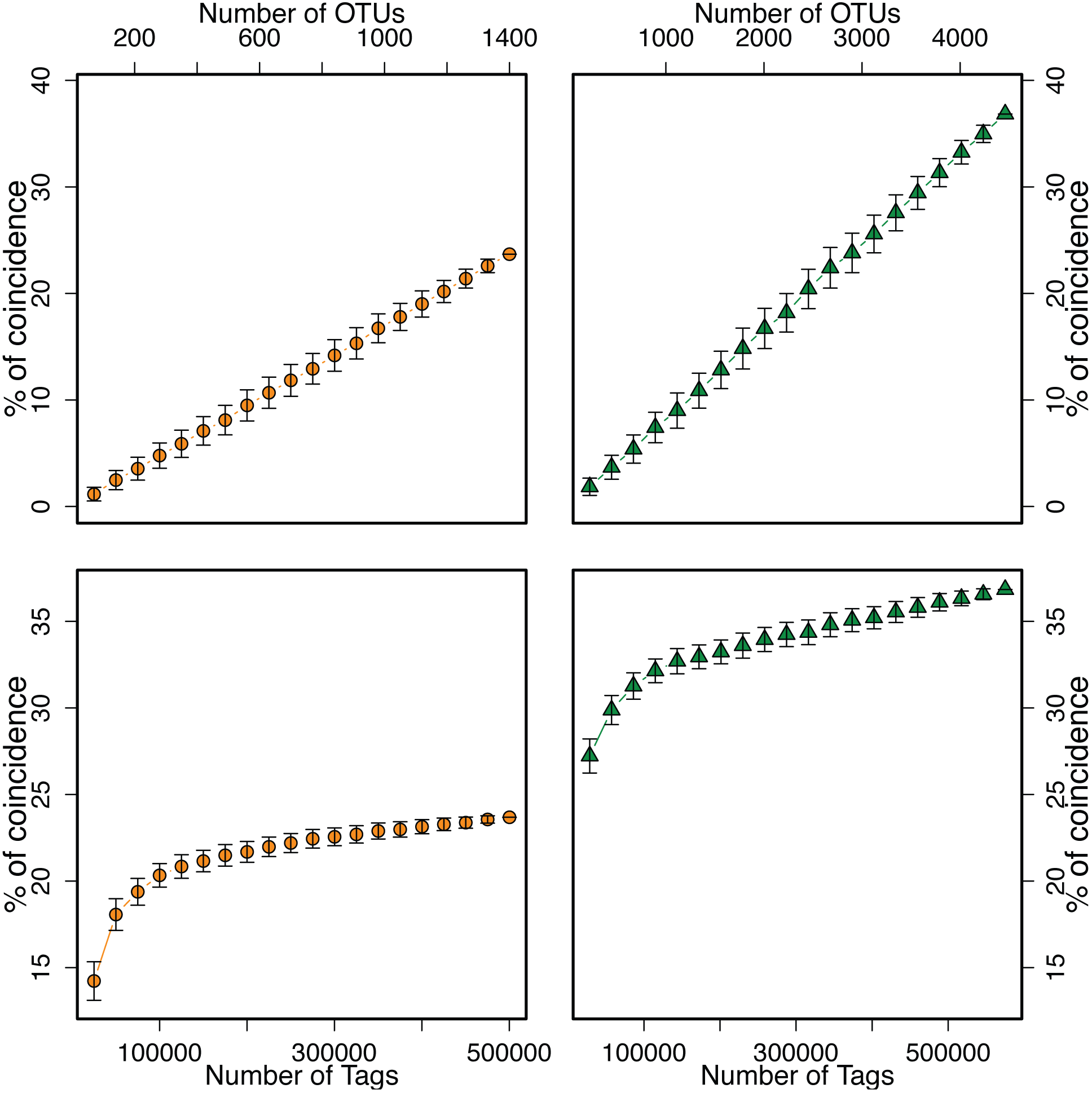
Accumulation curves of the percentage of cultured isolated found in the 454-pyrosequencing datasets (surface sample: A, C and bottom sample: B, D) when increasing the number of sampled OTUs (A, B) and Tags (C, D).

### 4. Comparisons across different times and locations

Three additional culture collections, isolated in the same way as the collection described above (see material and methods), were available for comparisons with 454-pyrosequencing data (A in Table 3). In addition to the 454-pyrosequecing dataset obtained in the present study, 454 sequences from the MODIVUS cruise in September 2007 were also available (see material and methods). These different datasets offered the possibility to compare how effective was the retrieval of isolated species in 454 sequence datasets at different time and space scales (B and C in Table 3). The number of cultures from each collection (columns 2 and 4 in Table 3) that could be compared to the two 454 datasets (B and C in Table 3) was different because the 16S rRNA fragments sequenced were different (see material and methods).

**Table 3.** Percentage of coincidence of the 454 pyrosequencing datasets from cruises SUMMER (September 2011) and MODIVUS (September 2007) with cultures covering the specific hypervariable regions sequenced in each cruise. The cultures belonged to four culture collections from the NW Mediterranean. The — symbol indicates that the comparison was not significant due to the low number of cultures with V1-V3 regions in the BBMO-3 culture collection. The comparison was Not Applicable (NA) when the samples compared did not belong to the same sampling site. The highest percentages of coincidences are indicated in bold type. BBMO means Blanes Bay Microbial Observatory.

| A) Culture collection | B) SUMMER 454 sequences<br>(September 2011) |  | C) MODIVUS 454 sequences<br>(September 2007) |  |  |  |  |
| --- | --- | --- | --- | --- | --- | --- | --- |
| 1 | 2 | 3 | 4 | 5 | 6<sup>b</sup> | 7<sup>b</sup> | 8<sup>b</sup> |
|  | # cultures<sup>a</sup> | % found with all sequences | # cultures<sup>a</sup> | % found with all sequences | % found with BBMO sequences | % found with Open-Sea, Surface | % found with Open-Sea, Bottom |
| BBMO-1 (2001-2004) | 65 | 8% | 39 | 44% | 23% | NA | NA |
| BBMO-2 (Feb 2007) | 11 | 9% | 20 | 80% | 40% | NA | NA |
| BBMO-3 (Sept 2007) | 2 | — | 6 | **83%** | **67%** | NA | NA |
| SUMMER (Sept 2011) | 38 | **37%** | 38 | 74% | NA | 26% | **34%** |

<sup>a</sup> Number of cultures in each collection where the sequence available matched the region targeted in the 454 sequencing (V1-V3 in Summer and V6 in MODIVUS).
<sup>b</sup> Comparison of the culture collections and the 454 pyrosequencing data of the sample collected at the same site as the sample for culture isolation.

Column 3 (Table 3) shows that the percent of cultures retrieved was much higher from the simultaneously taken culture collection (37%) than form those taken years before (~ 10%). Likewise, column 5 (Table 3) shows that the percent of cultures isolated the same year of the cruise MODIVUS was higher (83% and 80%) than with those isolated earlier (44% in 2001-2004) or later (74% in 2011). The cultures collected the same year but at a different date (February 2007) were retrieved almost with the same efficiency as those collected on the same dates (80%).

The next comparisons examine the efficiency of retrieval at different space scales. Columns 5 and 6 (Table 3) show the difference in percent of cultures found in the MODIVUS database if the whole set of sequences is considered (including a transect and a vertical profile, column 5 in Table 3) or if only the sequences from BBMO are used for the comparison (column 6 in Table 3). For the three BBMO collections the percent increased when a larger area was considered. The same can be observed in columns 5, 7 and 8 (Table 3) for the SUMMER culture collection: when only the sequences from the open sea station were included, the percentages recovered were 24 (for the surface sequences) and 34 (for the bottom sequences). When the whole MODIVUS dataset was included, 74% of the cultures were retrieved. Surprisingly, this percentage is larger than that of the SUMMER 454 dataset. This occurred despite the fact that the SUMMER data set had many more sequences (1.7×10^6^) than the MODIVUS dataset (3×10^5^).

## Discussion

### 1 Estimates of richness

A large proportion of the microbial world is invisible to traditional cultivation approaches but it can be accessed using molecular tools (DeLong, 1997). The first cloning and Sanger sequencing approaches revealed a wealth of new taxa in the oceans (Giovannoni *et al*., 1990). The number of clones examined however, was still too limited and there was a discrepancy between the taxa obtained in pure culture and by cloning and sequencing (Pedrós-Alió, 2006). The development of massive parallel sequencing technologies in the last decade has helped to uncover a large fraction of the hidden diversity of marine microorganisms (Sogin *et al*., 2006; Pedrós-Alió, 2012). In a previous study (Pommier *et al*., 2010) we used pyrosequencing of the V6 region of the 16S rDNA gene to estimate richness of the bacterial assemblages in the NW Mediterranean Sea. Around 20 000 tag sequences were obtained per sample. For the surface and deep samples from station D, we found 632 and 2 065 OTUs respectively and using the Chao estimate these translated into estimated richness of 1 289 and 4 156 OTUs for surface and deep samples respectively. It is well know that the number of new species retrieved increases with sampling size and sampling effort (Preston, 1960; Magurran, 1988; Rosenzweig, 1995) and a large part of the diversity remains hidden due to sampling limitations (Chao *et al*., 2009; Gotelli & Colwell 2011). Therefore, taking advantage of the increasing sampling depth of pyrosequencing, we obtained 1 million raw 16S rRNA gene tag-sequences per sample, trying to achieve realistic estimates of the whole bacterial diversity in our samples.

Our collector curves were close to reaching an asymptote (Figure 1). The relatively low percentage of singletons (~17%) found in this study compared to the higher percentage (40%-60%) in the previous study (Pommier *et al*., 2010) indicates a good coverage of the whole bacterial richness. By increasing the number of tags per sample from 20 000 to 1 000 000 we increased the number of OTUs from 632 to 1 400 at the surface and from 2 065 to 4 460 at the bottom. The Chao estimators were relatively close in both studies 1 289 vs 1 643 and 4 156 vs 5 029 for surface and bottom samples respectively. This suggests that the sequencing effort carried out in the present study is not necessary for a general evaluation of the richness of a marine sample, which is good news for further routine studies. However, we needed this effort to answer the questions discussed below.

The Chao index is based on the number of singletons and doubletons. We wanted to use all the information contained in the species accumulation curves to get a more robust estimate of richness. After testing several options, the Weibull cumulative function proved to be excellent. The extraordinary good fit (99% model efficiency) of this function to the species accumulation curves (Figure 2) gives confidence to the richness estimations made by extrapolation. Again, it is good for further studies that there were no significantly differences from the non-parametric estimations (z-tests on the differences, P > 0.05). The Weibull cumulative function has been satisfactorily fitted to species accumulation curves of well-sampled communities of birds (Flather, 1996), reptiles (Thompson *et al*., 2003), snakes (van Rooijen, 2012), butterflies (Jiménez-Valverde *et al*., 2006) and microplankton (Cermeño *et al*., 2014). We also fitted the model to a marine bacterial sample collected in the Western English Channel (Caporaso *et al*., 2012; Gibbons *et al*., 2013). This sample was sequenced at an extraordinary depth (on the order of 10^7^ Illumina tags) and we also found a good fit (R^2^=0.99). The fit was better when singletons were included in the analysis (the AIC value was lower, supplementary Fig. 1). Therefore, the Weibull cumulative function will be a very useful tool to characterize the species accumulation curves and to obtain robust richness estimates, at least for well-sampled bacterial communities.

The high abundance of the most abundant OTU in the surface sample (Figure 2) may have caused less OTUs to be uncovered, forcing the richness to appear lower at the surface sample than at the bottom sample. However, higher richness as well as higher diversity at the bottom than at the surface have been reported before for the sampling area (Pommier *et al*., 2010). Also, the high number of sequences examined and the richness estimations (Table 1) leave not doubt that a high percentage of the rare OTUs were sampled. Moreover, the number of OTUs in both samples was close to the number of OTUs estimated by other authors for the upper ocean (Rusch *et al*., 2007; Pommier *et al*., 2010; Crespo *et al*., 2013) and deep waters (Salazar *et al*., submitted).

The study of the English Channel mentioned above (Caporaso *et al*., 2012; Gibbons *et al*., 2013) is particularly relevant for our analysis. Station L4 was very deeply sequenced (10 million sequences) by Illumina. The sequences were then compared to pyrosequencing data from 72 samples collected along the seasons through six years (Caporaso *et al*., 2012) or to the ICoMM pyrosequencing samples from around the world’s oceans (Gibbons *et al*., 2013). The purpose of this exercise was to see whether the collectors curve could be saturated with sufficiently deep sequencing and whether the rare biosphere of one site, at one particular time, included all the taxa found at that site through the years. Caporaso *et al*. (2012) found that around 95% of the OTUs in the combined seasonal samples could be found in the one deeply sequenced sample. Thus, the conclusion was that the rare biosphere included all the taxa that were abundant at L4 at one time or another. “This suggested that the vast majority of taxa identified in this ecosystem were always present, but just in different proportions” (Caporaso *et al*., 2012).

However, there are some caveats in interpreting this data set. The number of OTUs found by pyrosequencing in the combined 72 samples was 13 424. We think this agrees well with our estimates of 2 000 to 5 000 OTUs in one single sample. A fourfold increase in OTUs when analyzing 72 samples collected through the seasons instead of 1 sample seems reasonable. On the other hand the deep Illumina sampling produced 116 107 OTUs, which is one order of magnitude higher.

Several issues must be considered. First, these authors found that 45–48% of their OTUs were singletons. Given than the data sets consisted of over 8×10^5^ sequences (seasonal samples) and the other of over 10^7^ sequences, these very large percentages of singletons are intriguing. In a previous study of the Mediterranean, we found a similar percentage of singletons (46%) with a lower number of sequences (≈2×10^5^) (Pommier *et al*., 2010; Crespo *et al*., 2013). However, with the increase in sequences to 5×10^5^ in the present study of the same area, the number of singletons was reduced to 17%. A reduction in percent of singletons with an increase in sequencing depth is what one would expect and is what other studies have found (Wall *et al*., 2009; Penton *et al*., 2013). Thus, we find the results of Caporaso *et al*. (2012) and Gibbons *et al*. (2013) surprising.

Second, most of the OTUs in Caporaso *et al*. (2012) could not have a taxonomy ascribed, even after exclusion of singletons. More than half were “unclassified bacteria” and about 10% were “unclassifiable (reads too short)” (see their Figure 1C). Thus, only about 40% of the OTUs could be classified. Again, this very low figure is intriguing. The short read length cannot be the reason, because in our earlier studies, we used exactly the same methodology and read length as for the seasonal samples from the English Channel or the world ocean samples from ICoMM and yet, 98.9% of our OTUs could be assigned a taxonomy at least at the Phylum level and most of the time at the Class level. Obviously, different criteria for assignment of identity can be responsible for this, but the 10% of sequences labeled as “unclassifiable (reads too short)” should probably have been discarded.

We used the data from their deep sequenced L4 sample to try the Weibull fit and found a good fit, both with and without the singletons. In both cases the number of OTUs was one order of magnitude higher than our estimates (40 000 and 100 000 OTUs without and with singletons respectively). It is difficult to integrate these very different estimates of richness. We see three possibilities. First, if both data sets and sequence processing are correct the English Channel could have ten times more species than the Mediterranean Sea. We find this possibility very unlikely, since both environments correspond to relatively open seawater. Second, perhaps the procedure chosen for OTU calling with the L4 data set overestimated the number of OTUs. As explained, the number of OTUs found in the seasonal samples was coherent with our estimates. The number of OTUs with the Illumina sequences, on the other hand was one order of magnitude higher. We believe that the current processing of pyrosequencing data is quite robust (Quince *et al*., 2011). Processing of Illumina tags, however, was still in its infancy when the studies mentioned above were carried out. Thus, the possibility exists of an overestimation of diversity. This is actually what happened in the first application of pyrosequencing to marine bacterial diversity in Sogin *et al*. (2006). Later studies found ways to properly clean the sequences and estimates became lower (Huse *et al*., 2010; Quince *et al*., 2011). We think this is the most likely explanation. Finally, there could be some methodological issue that caused an increase in diversity when the number of sequences increased by one order of magnitude. Thus, when diversity of marine bacterial communities was estimated from conventional clone libraries (with a few hundred clones) richness estimations were on the order of a few hundred OTUs. When similar samples were analyzed by HTS (with tens of thousands of sequences per sample) the richness estimators gave numbers of several thousand OTUs (as in the present study).

### 2. Comparison of sequencing and isolation

The current power of massive parallel sequencing allows probing the rare biosphere (Caporaso *et al*., 2012; Pedrós-Alió, 2012; Gibbons *et al*., 2013), but culturing is an alternative avenue to explore it (Pedrós-Alió, 2006, Shade *et al*., 2012). Comparing both approaches we have found that isolation retrieves some of the rarest taxa since only 24 to 37% of the isolates were found in the 454-pyrosequencing data and, moreover, they were found in extremely low abundance (Figure 3, Table 2).

In principle, the low percentage of coincidence might have been due to potential PCR bias and differential DNA amplification of the sequencing techniques (Berry *et al*., 2011; Pinto & Raskin, 2012). However, when tested *in silico*, the primers used for pyrosequencing covered the whole diversity captured by the primers used for Sanger sequencing of the isolates and therefore, a PCR bias affecting the diversity found using both methods was unlikely.

Our results showed that the deep sequencing approach failed to retrieve 76% of the bacterial cultures that were isolated in the surface sample. In a similar study, but with much shallower sequencing depth (~2 000 sequences per sample), Shade *et al*. (2012) found out that 61% of their isolates were not retrieved by 454-pyrosequencing. They performed pyrosequencing of a pool of bacterial cultures isolated from soil samples instead of Sanger sequencing of each isolate as we did in this study. Thus both studies, one in soils and another in seawater, produced similar percentages.

The long tail of rare species estimated by fitting the Weibull cumulative function to the accumulation curve (Figure 2, Table 1) and by the non-parametric Chao method (Chao *et al*., 2009) (Table 1) should harbor the isolates not found with the sequencing depth applied in this study. Increasing the number of tags sequenced would slowly uncover the isolated bacteria in a logarithmic way (Figure 4). Even if the whole bacterial diversity were found by increasing the sampling depth, culturing would still be essential for the study of marine bacterial communities, especially if the target is the rare biosphere (Donachie *et al*., 2007).

Taking advantage of other datasets previously collected from the same area of our sampling, we were able to compare 454-pyrosequecing data and cultured isolates from different times of the year and stations along a horizontal transect and a vertical profile (Table 3). The percentage of cultured isolates sequences found in the 454-pyrosequecing datasets decreased as the time between the collection of samples for isolation and pyrosequencing increased; i.e., cultures isolated from the same sample as the pyrosequencing data were found more easily in the pyrosequencing dataset than the cultures collected further away in time or space (Table 3). The consistency of this pattern within the two datasets, SUMMER and MODIVUS (Table 3), suggests that the composition of the bacteria isolated changes along the year probably following the annual succession of the marine bacterial composition (Alonso-Sáez *et al*., 2007; Alonso-Sáez *et al*., 2008; Gilbert *et al*., 2012). Similarly, Lekunberri *et al*. (2014) found that sequences from cultures obtained from one season of the year tended to cluster together with environmental sequences obtained at the same time of the year. These two findings suggest that the composition of rare taxa also follows an annual succession, supporting the conclusion that the environment determines the marine bacterial composition, not only of the abundant taxa but also of the rare members of the bacterial community (Galand *et al*., 2009). The shorter length of the 454-tag sequences (and the consequent decrease in precision of taxonomy) was likely responsible for the higher percentage of cultured isolates found on the 454-pyrosequecing datasets of MODIVUS cruise (454-tags sequences of ~68 bp) than of SUMMER cruise (454-tags sequences of ~400 bp).

In conclusion, by obtaining 10^6^ tags per sample and fitting a Weibull cumulative function we have been able to obtain a robust estimate of the richness of the bacterial assemblages in two samples at the surface and deep Mediterranean Sea. This richness is of the order of 3 000 to 5 000 taxa. The comparison with cultures shows that many of the isolates are found deep within the rare biosphere, and we have determined the sequencing effort necessary to retrieve them. HTS and culturing appear as two complementary strategies to probe the fabric of the rare biosphere.

## Acknowledgements

We thank the crews and scientists in cruises Modivus and SUMMER, both on the RV García del Cid, supported by the Spanish MICINN grants CTM2005-04795/MAR and CTM2008-03309/MAR respectively.

We thank F. M. Cornejo-Castillo for his advice on the method for isolates’ differentiation. E. Sa and V. Balagué help with bacterial culturing and PCR work is highly appreciated.

B. G. C. was supported by a Juan de la Cierva contract from the Spanish “Ministerio de Ciencia e Innovación”. Research was funded by the Spanish “Plan Nacional de Investigación Científica y Técnica” grants Marine Gems (CTM2010-20361) and Blue Genes (CTM2013-48292-C3-1-R).

## References

Alonso-Sáez L, Balagué V, Sá EL, Sánchez O, González JM, Pinhassi J, et al. (2007). Seasonality in bacterial diversity in north-west Mediterranean coastal waters: assessment through clone libraries, fingerprinting and FISH. FEMS Microbiol Ecol 60:98–112.

Alonso-Sáez L, Sánchez O, Gasol JM, Balagué V, Pedrós-Alio C. (2008). Winter-to-summer changes in the composition and single-cell activity of near-surface Arctic prokaryotes. Environ Microbiol 10:2444–54.

Amaral-Zettler L, Artigas LF, Baross J, Bharathi LPA, Boetius A, Chandramohan D, et al. (2010). A Global Census of Marine Microbes — Census of Marine Life Maps and Visualization. In:Life in the World’s Oceans: Diversity, Distribution, and Abundance., McIntyre., A (ed), Wiley-Blackwell, pp. 223–345.

Berry D, Ben Mahfoudh K, Wagner M, Loy A. (2011). Barcoded primers used in multiplex amplicon pyrosequencing bias amplification. Appl Environ Microbiol 77:7846–9.

Caporaso J, Kuczynski J, Stombaugh J, Bttinger K, Bushman F, Costello E, et al. (2010). QIIME allows analysis of high-throughput community sequencing data. Nature 7: 335–336.

Caporaso JG, Paszkiewicz K, Field D, Knight R, Gilbert J a. (2012). The Western English Channel contains a persistent microbial seed bank. ISME J 6:1089–93.

Cermeño P, Teixeira IG, Branco M, Figueiras FG, Marañón E. (2014). Sampling the limits of species richness in marine phytoplankton communities. J Plankton Res 36:1135–1139.

Chao A, Colwell RK, Lin C, Gotelli NJ. (2009). Sufficient Sampling for Asymptotic Minimum Species Richness Estimators. Ecology 90:1125–1133.

Crespo BG, Pommier T, Fernández-Gómez B, Pedrós-Alió C. (2013). Taxonomic composition of the particle-attached and free-living bacterial assemblages in the Northwest Mediterranean Sea analyzed by pyrosequencing of the 16S rRNA. Microbiologyopen 2:541–552.

Curtis TP, Sloan WT, Scannell JW. (2002). Estimating prokaryotic diversity and its limits. Proc Natl Acad Sci 99:10494–99.

DeLong E. (1997). Marine microbial diversity: the tip of the iceberg. Trends Biotechnol 15:203–207.

Donachie SP, Foster JS, Brown M V. (2007). Culture clash: challenging the dogma of microbial diversity. ISME J 1:97–9.

Dowd SE, Callaway TR, Wolcott RD, Sun Y, McKeehan T, Hagevoort RG, et al. (2008). Evaluation of the bacterial diversity in the feces of cattle using 16S rDNA bacterial tag-encoded FLX amplicon pyrosequencing (bTEFAP). BMC Microbiol 8:125.

Eilers H, Pernthaler J, Glöckner FO, Amann R. (2000). Culturability and In situ abundance of pelagic bacteria from the North Sea. Appl Environ Microbiol 66:3044–51.

Erwin T. (1991). How many species are there? Revisited. Conserv Biol 5:1–4.

Fisher MM, Triplett EW. (1999). Automated approach for ribosomal intergenic spacer analysis of microbial diversity and its application to freshwater bacterial communities. Appl Environ Microbiol 65:4630–6.

Flather CH. (1996). Fitting Species-Accumulation Functions and Assessing Regional Land Use Impacts on A vian Diversity Curtis H. Flather. J Biogeogr 23:155–168.

Galand PE, Casamayor EO, Kirchman DL, Lovejoy C. (2009). Ecology of the rare microbial biosphere of the Arctic Ocean. Proc Natl Acad Sci 106:22427–22432.

Gibbons SM, Caporaso JG, Pirrung M, Field D, Knight R, Gilbert JA. (2013). Evidence for a persistent microbial seed bank throughout the global ocean. Proc Natl Acad Sci 110:4651–4655.

Gilbert JA, Steele JA, Caporaso JG, Steinbrück L, Reeder J, Temperton B, et al. (2012). Defining seasonal marine microbial community dynamics. ISME J 6:298–308.

Giovannoni SJ, Britschgi TB, Moyer CL, Field KG. (1990). Genetic diversity in Sargasso Sea bacterioplankton. Nature 345:60–3.

Hagström Å, Pommier T, Rohwer F, Simu K, Svensson D, Zweifel U. (2002). Bioinformatics reveal surprisingly low species richness in marine bacterioplankton. Appl Environ Microbiol 67: 3628–3633.

Huse SM, Welch DM, Morrison HG, Sogin ML. (2010). Ironing out the wrinkles in the rare biosphere through improved OTU clustering. Environ Microbiol 12:1889–98.

Jiménez-Valverde A, Mendoza SJ, Cano JM, Munguira ML. (2006). Comparing Relative Model Fit of Several Species-Accumulation Functions to Local Papilionoidea and Hesperioidea Butterfly Inventories of Mediterranean Habitats. Biodivers Conserv 15:177–190.

Jones SE, Lennon JT. (2010). Dormancy contributes to the maintenance of microbial diversity. Proc Natl Acad Sci U S A 107:5881–6.

Kindt R, Coe R. (2005). Tree diversity analysis. A manual and software for common statistical methods for ecological and biodiversity studies. World Agroforestry Centre (ICRAF): Nairobi (Kenya).

Lekunberri I, Gasol JM, Acinas SG, Gómez-Consarnau L, Crespo BG, Casamayor EO, et al. (2014). The phylogenetic and ecological context of cultured and whole genome-sequenced planktonic bacteria from the coastal NW Mediterranean Sea. Syst Appl Microbiol. 37:216–28.

Lynch MDJ, Bartram AK, Neufeld JD. (2012). Targeted recovery of novel phylogenetic diversity from next-generation sequence data. ISME J 6:2067–77.

Magurran AE. (1988). Ecological diversity and its measurements. Princeton University Press: Princeton, New Jersey.

May RM. (1988). How many species are there on Earth? Science 241:1441–9.

Mora C, Tittensor DP, Adl S, Simpson AGB, Worm B. (2011). How many species are there on Earth and in the ocean? PLoS Biol 9:e1001127.

Oksanen J, Guillaume-Blanchet F, Kindt R, Legendre P, Minchin P, O’Hara R, et al. (2013). Vegan: Community Ecology Package.

Pace NR. (1997). A molecular view of microbial diversity and the biosphere. Science 276:734–40.

Pedrós-Alió C. (2006). Marine microbial diversity: can it be determined? Trends Microbiol 14:257–63.

Pedrós-Alió C. (2012). The Rare Bacterial Biosphere. Ann Rev Mar Sci 4:449–466.

Pedrós-Alió C, Calderón-Paz J-I, Guixa-Boixereu N, Estrada M, Gasol JM. (1999). Bacterioplankton and phytoplankton biomass and production during summer stratification in the northwestern Mediterranean Sea. Deep Sea Res Part I Oceanogr Res Pap 46:985–1019.

Penton CR, St Louis D, Cole JR, Luo Y, Wu L, Schuur EAG, et al. (2013). Fungal diversity in permafrost and tallgrass prairie soils under experimental warming conditions. Appl Environ Microbiol 79:7063–72.

Pinto AJ, Raskin L. (2012). PCR biases distort bacterial and archaeal community structure in pyrosequencing datasets. PLoS One 7:e43093.

Pommier T, Neal P, Gasol J, Coll M, Acinas S, Pedrós-Alió C. (2010). Spatial patterns of bacterial richness and evenness in the NW Mediterranean Sea explored by pyrosequencing of the 16S rRNA. Aquat Microb Ecol 61:221–233.

Preston FW. (1960). Time and Space and the Variation of Species. Ecology 41:612–627.

Quast C, Pruesse E, Yilmaz P, Gerken J, Schweer T, Yarza P, et al. (2013). The SILVA ribosomal RNA gene database project: improved data processing and web-based tools. Nucleic Acids Res 41:D590–6.

Quince C, Lanzen A, Davenport RJ, Turnbaugh PJ. (2011). Removing noise from pyrosequenced amplicons. BMC Bioinformatics 12:38.

R CoreTeam. (2013). R: A language and environment for statistical computing.

Ritz C, Streibig J. (2005). Bioassay Analysis using R. J Stat Softw 12.

Van Rooijen J. (2012). Estimating the snake species richness of the Santubong Peninsula (Borneo): a computer-simulation. Amphibia-Reptilia 33:521–525.

Rosenzweig M. (1995). Species diversity in space and time. In:Cambridge University Press: Cambridge.

Rusch DB, Halpern AL, Sutton G, Heidelberg KB, Williamson S, Yooseph S, et al. (2007). The Sorcerer II Global Ocean Sampling expedition: northwest Atlantic through eastern tropical Pacific. PLoS Biol 5:e77.

Schauer M, Balagué V, Pedrós-Alió C, Massana R. (2003). Seasonal changes in the taxonomic composition of bacterioplankton in a coastal oligotrophic system. Aquat Microb Ecol 31:163–174.

Scheinert P, Krausse R, Ullmann U, Söller R, Krupp G. (1996). Molecular differentiation of bacteria by PCR amplification of the 16S–23S rRNA spacer. J Microbiol Methods 26:103–117.

Schloss PD, Gevers D, Westcott SL. (2011). Reducing the effects of PCR amplification and sequencing artifacts on 16S rRNA-based studies. Gilbert, JA (ed). PLoS One 6:e27310.

Shade A, Hogan CS, Klimowicz AK, Linske M, McManus PS, Handelsman J. (2012). Culturing captures members of the soil rare biosphere. Environ Microbiol 14:2247–52.

Sogin ML, Morrison HG, Huber JA, Mark Welch D, Huse SM, Neal PR, et al. (2006). Microbial diversity in the deep sea and the underexplored “rare biosphere”. Proc Natl Acad Sci USA 103:12115–20.

Staley J, Konopka A. (1985). Measurement of in situ activities of nonphotosynthetic microroganisms in aquatic and terrestrial habitats. Annu Rev Microbiol 39:321–383.

Thompson GG, Withers PC, Pianka ER, Thompson S a. (2003). Assessing biodiversity with species accumulation curves; inventories of small reptiles by pit-trapping in Western Australia. Austral Ecol 28:361–383.

Wall PK, Leebens-Mack J, Chanderbali AS, Barakat A, Wolcott E, Liang H, et al. (2009). Comparison of next generation sequencing technologies for transcriptome characterization. BMC Genomics 10:347.

